# IKZF2-ERBB4 gene fusion leads to *ERBB4* overexpression in Anal Squamous Cell Carcinoma but with no benefit for ERBB4 inhibition

**DOI:** 10.1101/2024.11.28.625910

**Authors:** Richon Sophie, Schnitzler Anne, Lazartigues Julien, Briaux Adrien, Vacher Sophie, Rania El Botty, El Alam Elsy, Mariani Pascale, Neuzillet Cindy, Lièvre Astrid, Cacheux Wulfran, Bièche Ivan, Dangles-Marie Virginie

## Abstract

Anal squamous cell carcinoma (ASCC) is a rare tumour, but with increasing incidence for local and metastatic tumours. Current therapy is based on chemoradiotherapy, associated with immunotherapy in advanced stages, with frequent side effects, poor results in advanced stages and recurrence. New therapeutics, including targeted therapies, are then needed. In this context, we identified here a *IKZF2*-*ERBB4* gene fusion in a patient ASCC tumour, leading to overexpressed mRNA encoding a functional ERBB4 protein. This gene fusion has been already reported in other tumour types but not yet investigated as therapeutical target. The matched patient tumour-derived xenograft displays the same gene fusion and mRNA overexpression. It provided biological material for anti-ERBB4 testing in different cell models (*in vivo* xenograft, *in vitro* cell cultures). We used afatinib and lapatinib, 2 chemical pan-ERBB inhibitors approved in clinics, with anti-ERBB4 properties. ERBB4 inhibition did not lead to tumour growth inhibition although afatinib and lapatinib dramatically decreased ERBB4 phosphorylation with impact on downstream signalling MAPK/ERK but not PI3K/AKT pathways. Likewise, specific *ERBB4* knock-outing did not affect tumour cell proliferation. These ‘negative’ results must be put in line with reported potential crosstalk between PI3K/AKT and MAPK/ERK pathways in therapeutic resistance.

## Introduction

Anal squamous cell carcinoma (ASCC) is a rare tumour, but with increasing incidence. The present standard of care still remains primary chemoradiotherapy, which results in a high level of disease control for small, early-stage ASCC. More advanced cancers poorly respond to this treatment, and the disease relapses locoregionally in 30% to 60% of patients, resulting in an abdominoperineal resection. New treatment strategies, including molecular targeted therapies are warranted in order to improve disease control while none of which have yet been approved for ASCC treatment [Rao et al, 2021; Laxmi et al, 2023]. For this purpose, a better understanding of molecular mechanisms involved in anal carcinogenesis might lead to the identification of new therapeutic targets as well as prognostic and predictive biomarkers. While ASCC was still ignored within The Cancer Genome Atlas program, some genomic studies depict the molecular landscape of ASCC [Bernardi et al, 2015; Chung et al, 2016; Morris et al, 2017; Cacheux et al, 2018; Cacheux et al, 2019; Necchi et al, 2022; Hamza et al, 2024]. We and others have then demonstrated the high importance of the network defined by phosphatidylinositol-3-kinase (PI3K), AKT and mammalian target of rapamycin (mTOR) pathways downstream of Receptor Tyrosine Kinase (RTK) in ASCC.

*ERBB4* encodes a member of the RTK class I family named ‘ErbB family’, a group of targetable receptor tyrosine kinases that includes Epidermal Growth Factor Receptor (*ERBB1*), HER2 (*ERBB2*), and ErbB3 (*ERBB3*). Besides, data support the relevance of targeting ErbB4 in non-small cell lung cancer and in head and neck squamous carcinoma [Kurppa et al, 2016; Nakamura et al, 2016; Zilberg et al, 2018].

We identify here *ERBB4* among RTKs as the most overexpressed RTK in ASCC and in particular by a gene fusion mechanism of activation. This *IKZF2-ERBB4* gene fusion and related *ERBB4* overexpression misled to higher sensitivity to ErbB4 inhibition, which we unfortunately did not find in Erbb4-overexpressing cells.

## Material and methods

### Patient samples

Human samples used here for molecular analysis have been previously described [Cacheux et al, 2019]. Briefly, these tumours have been collected from patients with ASCC treated between 1992 and 2015 at the Institut Curie Hospital Supporting Information **Supplemental Table S1**). Tumour residual specimens were macrodissected to achieve maximum tumour purity (minimum 70%). Fifteen samples of adjacent normal anal squamous cell tissue from patients with ASCC were used as sources of normal RNA for RT-qPCR. Tissues samples were stored at −70°C until DNA and RNA extractions. This retrospective study was reviewed and approved by the Institut Curie Ethics Committee (No. A10-024).

### Patient tumour-derived xenograft

The Patient tumour-derived xenograft (PDX) model PdxAC17 was established and routinely passaged by subcutaneous engraftment with Matrigel in interscapular fat pad (BD Bioscience) in 5 to 7 week-old female RjOrl:NMRI-Foxn1^nu^/Foxn1^nu^ mice (Janvier Laboratories, Le Genest St Isle, France) using protocols and animal housing in accordance with national and European regulations and international guidelines (Project authorization: 02163.02).

### Genomic DNA extraction

The Qiagen DNeasy Tissue kit and the protocols for fresh frozen ASCC tissues were used. DNA was purified by column purification with a filter membrane and stored at −20°C before use.

### Array-CGH

Array-CGH experiments were carried out using 400K human genome CGH microarray and standard Agilent protocols (Agilent Technologies, Santa Clara, CA). Commercial human genomic DNA (Agilent Technologies) was used as diploid reference. Briefly, 1-1.5 μg of reference DNA and the same amounts of tumour DNA were digested with Alu1 and Rsa1 (Promega, Madison, WI, USA). The digested reference DNA fragments were labelled with cyanine 3-dUTP, and tumour DNA was labelled with cyanine 5-dUTP (Agilent Technologies). After clean-up, labelled reference and tumour DNA were mixed as probes and hybridized onto an Agilent 400K human genome CGH microarray (Agilent Technologies) for 40 hours. Washing, scanning, and data extraction procedures were carried out according to standard protocols. Data were extracted using Feature Extraction software (v11.1), and normalized data were analyzed and visualized by Agilent Cytogenomics Edition 2.9.2.4 (Agilent Technologies). Agilent Cytogenomics v4.0.3 was used to calculate the log2 ratio for every probe to identify genomic aberrations. Only genomic aberrations longer than three consecutive probes with an abnormal log2 (log2 ratio > 1.0 or < -1.0) ratio were taken into account for analysis. The mean log2 ratio of all probes in a chromosome region between 0.30 and 1.0 was classified as genomic gain, more than 1.0 (with a size < 10Mb) as focal amplification, less than -0.30 as heterozygous deletion and less than -1.0 (with a size <5Mb) as homozygous deletion.

### Total RNA extraction

Total RNA was extracted from fresh frozen ASCC and normal anal tissue by the acid-phenol guanidium method and quantified using an ND-1000 NanoDrop Spectrophotometer with its corresponding software (Thermo Fisher Scientific Inc., Wilmington, DE). RNA quality was determined by electrophoresis on agarose gel with ethidium bromide staining. The 18S and 28S RNA bands were visualized under ultraviolet light. Total RNA was stored at −20°C before use.

### qRT-PCR

Quantitative RT-PCR was conducted on RNA extract from tumour samples. Quantitative values were obtained from the cycle number (Ct value) at which the increase in the fluorescence signal associated with exponential growth of PCR products started to be detected by the laser detector of the ABI Prism 7900 Sequence Detection System (Perkin-Elmer Applied Biosystems, Foster City, CA), using PE Biosystems analysis software according to the manufacturer’s manuals. Gene primers were designed with the assistance of Oligo 6.0 software (National Biosciences, MN). All nucleotide sequences are shown in **Supplemental Table S2A.** Each sample was normalized based on its *TBP* content (Genbank accession NM_003194). Results, expressed as N-fold differences in target gene expression relative to the TBP gene and termed “Ntarget,” were determined as Ntarget = 2^ΔCt^ sample, where the ΔCt value of the sample is determined by subtracting the average Ct value of the target gene from the average Ct value of the TBP gene. The Ntarget values of the samples were subsequently normalised such that the Ntarget value median of the 15 adjacent normal anal tissues was equal to 1.

### Fusion Transcript Characterization

To detect fusion transcripts, we design the forward primer targeting the 5’ partner gene and reverse primer targeting the 3’ partner. Primer pairs (see **Supplemental Table S2B** for sequences) for the coding exons of the fusion genes were generated using Oligo 4.0. PCR products were separated by gel electrophoresis in a 3% agarose gel and visualized by ethidium bromide staining. Primers were also designed to span the predicted exons forming the fusion breakpoint. After cDNA synthesis from total RNA extracts using Reverse Transcription Kit, according to the manufacturer’s protocol (QIAGEN, GmbH), IKZF2-ERBB4 transcript fragments were obtained by PCR amplification. The amplified products were sequenced with a BigDye Terminator kit on an ABI Prism 3130 automatic DNA sequencer (Applied Biosystems, Courtabæuf, France). The sequences were compared with the corresponding cDNA reference sequence (IKZF2: NM_001079526; ERBB4: NM_005235.2) to characterize the fusion transcript.

Using the NCBI ORF Finder (http://www.ncbi.nlm.nih.gov/gorf/gorf.html), the gDNA sequence of 12kb was analyzed to check the presence of open reading frame and identify the start codon.

### Cell proliferation assays and chemical blocking

Short-term primary culture of AC17 cells has been established from PDXAC17 tissue. For this purpose, PDXAC17 tissue, was first finely cut with scalpel blade and crushed. The resulting pieces were transferred into a 25 cm^2^ culture flask in ‘culture medium’ (DMEM supplemented with 10% heat-inactivated fetal calf serum (Invitrogen, France), penicillin, streptomycin and amphotericin (Anti-Anti 100x, Gibco, France) and incubated at 37°C, 5% CO2. Floating canalospheres were observed within 48h as previously reported in colon cancer [Weiswald et al, 2013] and transferred into a 25 cm^2^ culture flask. Maintained on tissue-culture-treated plastic, floating cancer spheres finally formed adherent cell sheets. After expansion though several passages, AC17 cells (passage P3) were collected using trypsinization for WST and Xcelligence assays. AC17 cell viability was assayed using the WST-1 kit (Roche), according to supplier’s instructions: 15,000 cells were plated in 96-well culture plates. Inhibitors or control medium were added 4 days later and incubated 3 days with cells. The following formula was used to calculate from absorbance data the relative proliferation index (%) = 100*(Asample –Ablanck)/(Acontrol-Ablanck). Real-time monitoring of cell proliferation studies was also performed on AC17 cells using xCELLigence system (Roche Inc., France) according to supplier’s instructions. AC17 cells were seeded at a density of 15,000 cells/well in 96-well E-Plates (ACEA Biosciences) and afatinib was added 30h later for real-time measuring of electrical impedance thanks to microelectrodes at the well bottom. Cell index, proportional to cell number, was registered every 15 minutes.

For inhibitor assays, afatinib (BIBW2992, Euromedex, France) was dissolved in DMSO and kept at – 20 °C before use. One lapatinib pill (250 mg, TYVERB, Novartis, France) was dissolved in MCT (0.5% methylcellulose, 0.2% Tween 80) by mixing using vortex for 1 minute.

### Western blot

Afatinib and lapatinib treatment was done on primary cancer cells. For that, PDXAC17 tissue sample was first finely cut with scalpel blade and crushed. The resulting pieces were transferred into a 25 cm^2^ culture flask in ‘culture medium’ and incubated at 37°C, 5% CO2. Floating canalospheres were transferred into a 25 cm^2^ culture flask and formed adherent cell sheets. Four days later, adherent cells were collected using trypsin and 2.75 10^6^ cells were seeded by well in a 6-well plate. The next day, anti-ERBB4 drugs were added and incubated for 3h. Protein lysates were resolved on 4–12% TGX gels (Bio-Rad®, France), transferred onto nitrocellulose membranes (Bio-Rad®) and immunoblotted with rabbit antibodies from Cell Signaling Technology (France) against GAPDH (#2118), AKT (#9272), P-AKT ((#4058, Ser473), MEK1/2 (#9126) and P-MEK1/2 (#9154) ; and from Epitomics (Burlingame, CA) against ERBB4 (#1200-1) and P-ERBB4 (#2295-1). After washes, membranes were incubated with the appropriate horseradish peroxidase-conjugated affinity-purified goat anti–rabbit secondary antibodies (Jackson ImmunoResearch Laboratories, UK). Quantification of protein expression was performed using Multi Gauge software and normalized to GAPDH expression. For each condition, the ratio of p-protein/protein was calculated.

### SiRNA inhibition

For siRNA-mediated *ERBB4* knockdown, viability of immortalized AC17 cell line was measured with Incucyte® approach. For that, adherent cells from canalospheres were collected using trypsin Cells were seeded in 1 ml in 6-well plates at a density of 1.5 10^6^ cells per well for incubation with 2ml of hTErt lentivirus (pLV-hTerT-IRES-Hygro, Addgène) and 50µl of polybrene 100X (Sigma) for 72h. After washing, cells were expanded through trypsin passages. Immortalized AC17 cells were subsequently infected with H2BRFP lentivirus (kindly provided by Eva Paluch Laboratory for Molecular cell Biology, UC London, England) as follows: 3.10^6^ cells were seeded in 3 ml in 6-well plates and incubated with 2ml H2BRFP lentivirus and 50µl polybrène 100X for 72h.

siRNAs against human ERBB4 (Hs_ERBB4_5 FlexiTube siRNA, #SI02223893 and Hs_ERBB4_6 FlexiTube siRNA, #SI02223900, Qiagen) or non-targeting control (NTC) (#SI03650325, Qiagen). Cells were plated in 6-well plates at a density of 2.10^5^ cells per well. siRNAs were added 24 hours after seeding at the final concentration of 25 nM with 12 to 24 μL HiPerFect (Qiagen). On day 5 after transfection, cells were collected for RNA.

Cells were seeded at 20,000 cells/well in 96-well culture plates in 100 µL ‘culture medium’. The plate was inserted into an Incucyte® Live-Cell Analysis System (Sartorius, Göttingen, Germany) for real-time imaging, with two fields imaged per well at 10x magnification every 2 h for a total of 4 days. The fluorescence and confluence data were analyzed using IncuCyte® Confluence version 2021 C software. The data shown represent fluorescence normalized to confluence at each time point.

### Histological analysis

Paraffin-embedded tissue sections (3 μm) were deparaffinized and rehydrated by a series of xylene and ethanol washes for HES staining. All histological examinations were performed under blind conditions (i.e. with no prior knowledge of the treatment).

### RNA *in situ* hybridization

*In situ* visualization of gene fusion transcript was done using BaseScope-ISH Assay protocol. For that, PdxAC17 was fixed in freshly prepared 4% PFA for 24 hours at 4°C. Then tissue was immersed in a 10% sucrose solution in 1X PBS then in 20% sucrose solution and finally in 30% sucrose solution until the tissue sinks to the bottom, all steps done at 4°C. Tissue was freezing in OCT (Optimal Cutting Temperature) embedding media with crushed dry ice. The block was sectioning to a thickness of 10µm and mounting on Superfrost Plus Slides. Tissue was washed with 1X PBS 5 min while moving the rack up and down to remove OCT, baked 30min at 60°c and post-fixed by immersing them in 4%PFA in 1X PBS. Tissue was dehydrated in an ascending series of ethanol 5 min each at room temperature. BaseScope Chromogenic Assay was perform by using BaseScope Duplex Detection Reagent kit (ACD Biotechne, ref 322381) with BaseScope Duplex Positive Control Probe-Human (Hs)- C1-Hs-PPIB-1ZZ/C2-POLR2A-1ZZ (ref 700101) and BaseScope Duplex Negative Control Probe-C1-DapB-1ZZ/C2-DapB-1ZZ (ref 700141) and specific BaseScope Probe BA-Hs-IKZF-ERBB4-Junc-C1 (ref 1245201-C1)/Hs-ERBB4-No-XMm-1zz-st- C2 (ref 1245211-C2). Dots was image on LIPSI microscope (Nikon Center, Institut Curie).

### Statistical analysis

If applicable, data are presented as the mean ± S.E.M. Tests for significant differences between the groups were performed using a Mann-Whitney U test using GraphPad Prism 5.0 (GraphPad software, San Diego, CA). A minimum *p* value of 0.05 was chosen as the significance level.

## Results and Discussion

### *ERBB4* as the most overexpressed gene encoding RTK in our ASCC cohort

Alterations of RTK genes are frequent in human cancers. RTK overexpression leads to increased local concentration of receptors, which results in elevated RTK signaling and overcomes the antagonizing regulatory effects [Du and Lovly, 2018]. Squamous cell carcinomas are also linked to frequent RTK gene amplifications, contributing to elevated receptor activity and potential poor prognosis [Dotto and Rustgi, 2016].

Thus, we tested 18 key RTK genes linked to cancer-targeted therapies (listed in **Supplemental Table S3**) in our 40 ASCC cohort by qRT-PCR and identified *ERBB4* as the gene whose expression could be the most increased compared to its expression in normal tissue (**Figure 1A**). Three out of the 40 ASCC specimens displayed a marked *ERBB4* overexpression with a 9 to 16-fold increase relative to the normal tissue median values (**Figure 1B**).

**Figure 1.**
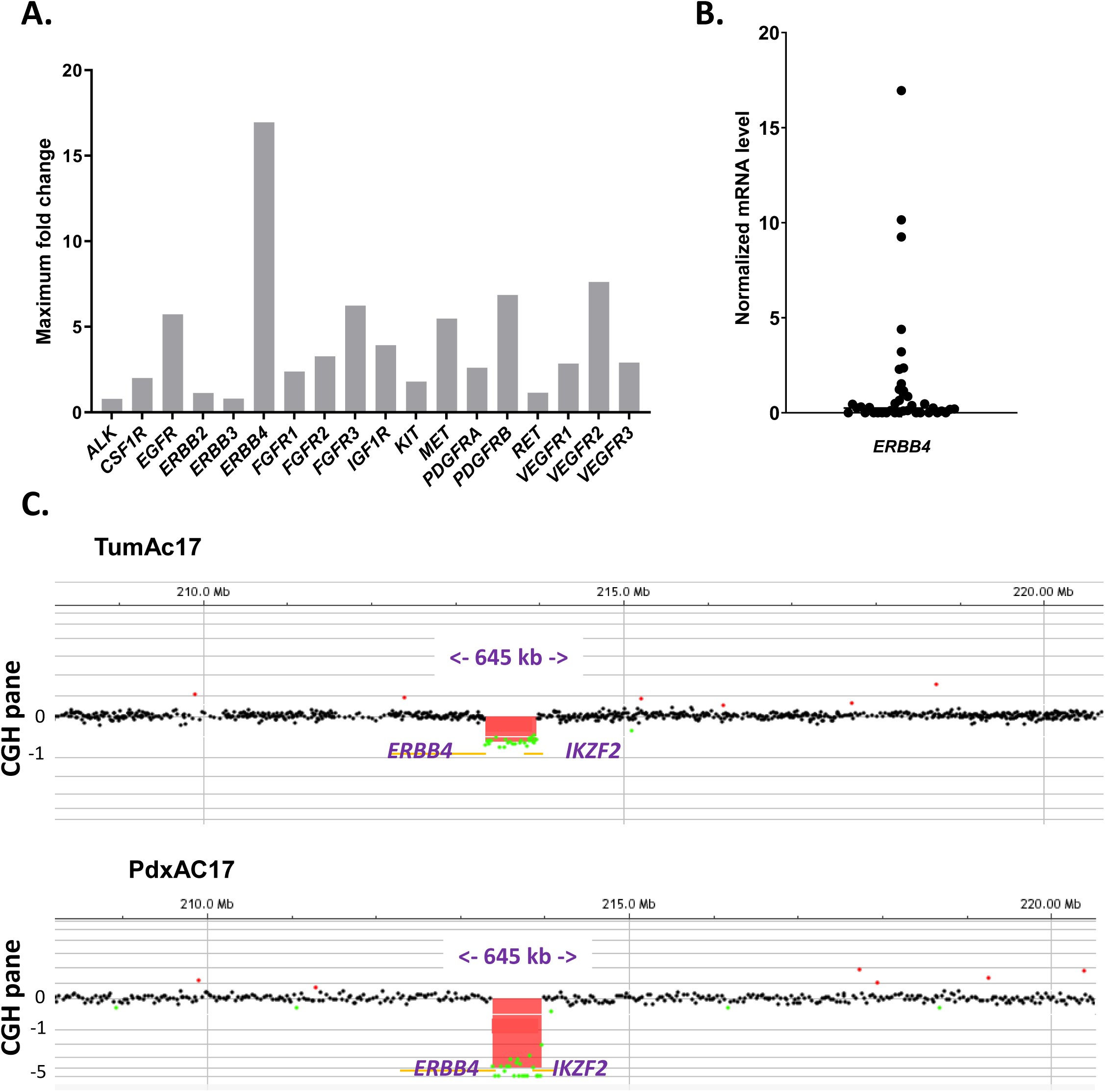
*ERBB4* as the most overexpressed gene encoding RTK in ASCC due to gene fusion. **A.** Maximum fold-change of mRNA expression levels observed in the 40 ASCC samples for each of the 18 studied genes encoding RTK, in comparison with its expression in normal tissue. **B.** *ERBB4* mRNA expressions in the 40 ASCC samples. As described in Materials and Methods section, each gene expression for each tumour sample has been doubly normalized in relation to its expression of *TBP* house-keeping gene and the median value in the 15 normal anal tissues. **C.** Genome-wide array CGH identified the same deletions encompassing the IKZF2 and ERBB4 genes in both patient tumour TumAC17 and derived PDX PdxAC17.

### *IKZF2-ERBB4* gene fusion involved in present *ERBB4* overexpression

As gene amplification is one of the basic mechanisms that lead to overexpression of oncogenes, we examined the amplification status of *ERBB4* locus by CGH in the 3 *ERBB4*- overexpressed tumour samples. No *ERBB4* gene amplification was observed. By contrast, among the three *ERBB4* overexpressing tumours, a homozygous focal intrachromosomal deletion of 645 Kb size was noted in TumAC17 (ERBB4 fold change≍ 12) encompassing *ERBB4* and *IKZF2* genes (**Figure 1C**). As nearly always reported for homozygous deletions [Priestley et al, 2019], this homozygous deletion is restricted to small chromosomal regions. These findings were suggestive of a rearrangement event leading to a potential *IKZF2-ERBB4* fusion transcript.

In order to characterize more deeply this genetic abnormality, we took advantage of TumAC17-derived xenograft we have established into immunodeficient mice to provide large quantity of human ASCC tissue and preclinical models. Firstly, we confirmed in the resulting PDX, referred as to PdxAC17, the presence of the same *IKZF2* and *ERBB4* homozygous gene deletion at the DNA level by CGH analysis (**Figure 1C**), confirming the interest of PDX to mimic the molecular profile of the origin tumour [Byrne et al, 2017]. Then, we used the RNA from both PDX and patient tumour tissue to confirm the absence of wild-type *ERBB4* transcript and the presence of the fusion transcript by RT-PCR with dedicated primers (**Figure 2A**). Like PCR assays, *in situ* hybridization confirmed (**Supplemental Figure S1**) the presence of *ERBB4* transcript only in fusion form. The *IKZF2e4-ERBB4e2* amplification yielded a PCR product of ∼500 pb, which was subsequently sequenced by Sanger method. Sanger sequencing confirmed that the junction of the two genes included exon 3 of *IKZF2* followed by exon 2 of *ERBB4* (**Figure 2B**). The same results were found in the corresponding patient tumour. The analysis of the cDNA sequence using NCBI ORF Finder suggested the presence of new open reading frame and identical start codon from IKZF2 protein. Thus, the in-frame gene fusion, with exons 1 to 3 of *IKZF2*, co-opts the *IKZF2* promoter for expression and retains all ERBB4 functional domains, including the ERBB4 intracellular domain (**Figure 2B**).

**Figure 2.**
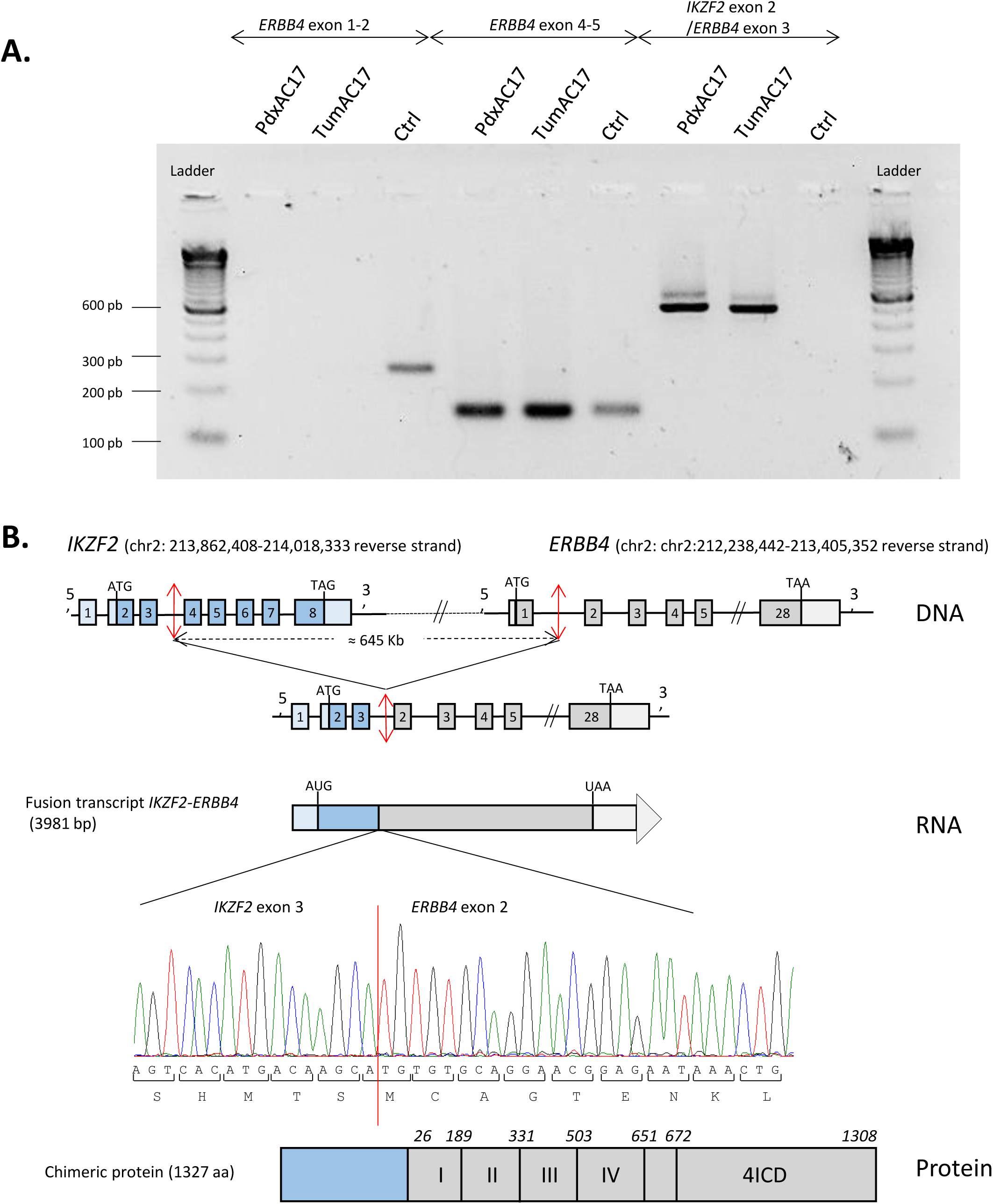
Identification of *IKZF2-ERBB4* fusion in TumAC17. **A.** PCRs with genomic DNA of TumAC17 and derived PdxAC17 were performed with 3 primer couples involving *ERBB4*. **B.** Sanger sequencing of the fusion site in *IKZF2-ERBB4* fusion. Schematic diagrams of gene rearrangements and ERBB4 protein domains in resulting IKZF2-ERBB4 fusion protein.

### Overview of the *IKZF2-ERBB4* gene fusion

The present gene fusion of *IKZF2-ERBB4* has been already described in one case of peripheral T cell lymphoma [Boddicker et al, 2016], one ovarian cancer cell line [Papp et al, 2018] and one non-small cell lung carcinoma [Priestley et al, 2019] associated with *ERBB4* overexpression for lymphoma and ovarian cases, while this kinase fusion is not yet reported in solid tumours in TCGA cohorts [Stransky et al, 2014; Gao et al, 2018]. More generally, *IKZF2-ERBB4* fusions have been reported in 20 tumours of Caris cohort (n=64,354) as the most common *ERBB4* fusion [Schubert et al, 2023]. This finding has to be put in line with a recent genomic analysis on 1,188 pancancer samples that has shown that more than 95% of gene fusions (3148 out of 3297) occurs in a unique tumour sample while recurrent gene fusions were often (59%; 46 out of 78 events) found across several histotypes like the present *IKZF2-ERBB4* fusion [PCAWG Transcriptome Core Group, 2020]. As expected, this fusion involving oncogene has been reported to tend to exhibit increased expression, by contrast to fusions involving tumour suppressors with decreased expression to samples without fusions [Gao et al, 2018].

To further investigate the impact of this gene fusion, we analysed IKZF2 and ERBB4 expression at protein level in different cell and cancer types using the Human Protein Atlas [Ponten et al, 2008, https://www.proteinatlas.org/]. We first focused our analysis on normal squamous epithelial cells to get an overview of expression level for IKZF2 and ERBB4 in cell lineages involved in squamous carcinoma. IKZF2 was found at high level in squamous cells in different locations while ERBB4 is poorly expressed (**Figure 3A**). The same analysis in different cancer types, including squamous carcinoma, shows consistent profiles: IKZF2 is highly expressed in various type of cancers by contrast to ERBB4 (**Figure 3B-C**), underlining the benefit for *ERBB4* of using *IKZF2* promoter within this fusion.

**Figure 3.**
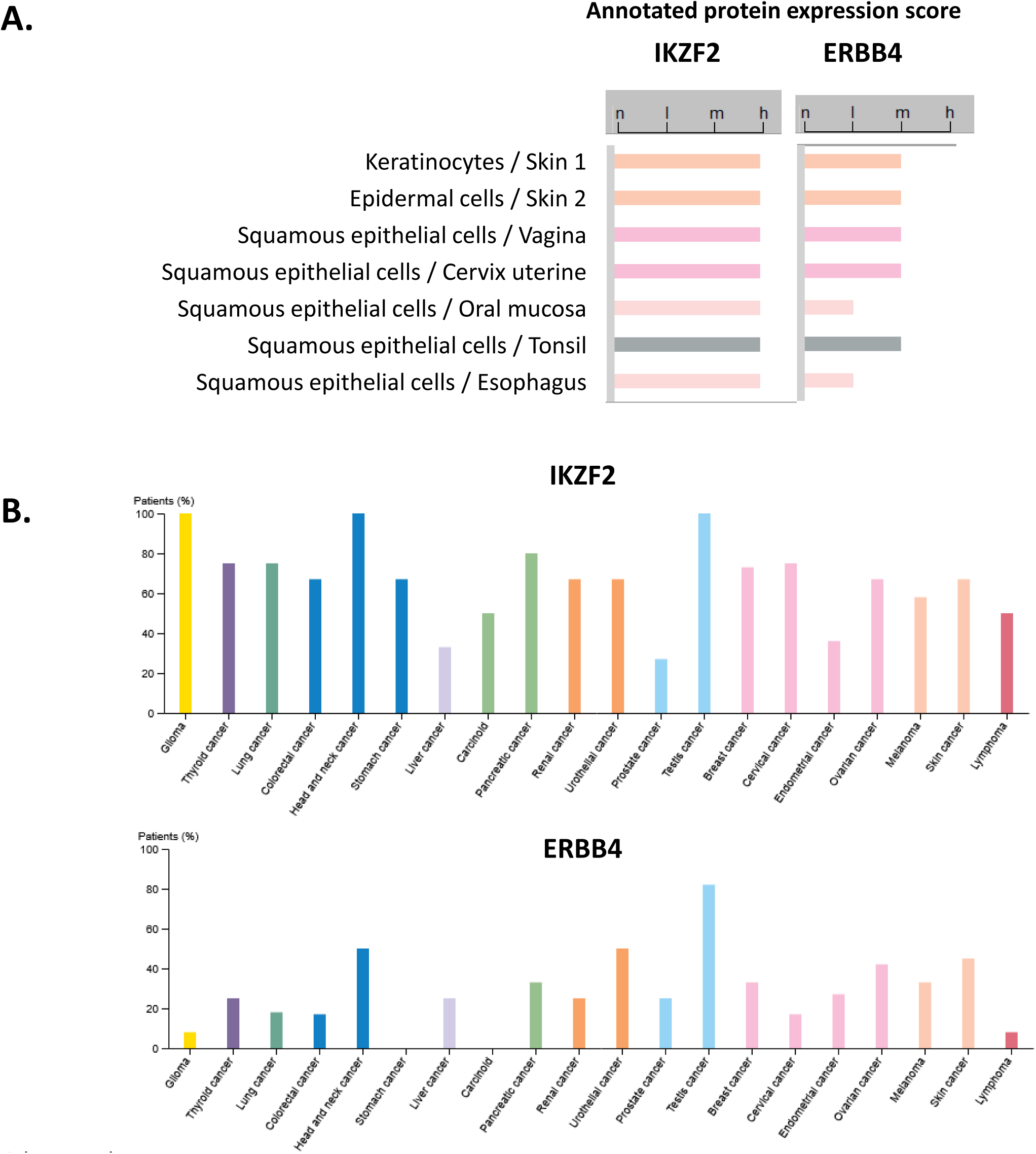
IKZF2 is expressed as a higher level than ERBB4 in squamous epithelial cells. **A.** According to THPA database, all studied normal squamous epithelial cells highly express IKZF2. By contrast, ERBB4 is moderately even poorly expressed in oral mucosa or esophagus. Annotated protein expression score is given in THPA as describing a knowledge-based best estimate of the protein expression in the annotated cell types as not detected (n), low (l), medium (m) or high (h), resulting of stringent evaluation of immunohistochemical staining pattern, RNA data from internal and external sources and available protein/gene characterization data. **B.** As for cancer tissues, bars indicate the percentage of patients (maximum 12 patients) with high and medium protein expression level. Low or not detected protein expression results in a white bar. Source for all presented histobars: https://www.proteinatlas.org/.

### Lack of sensitivity to anti-ErbB4 inhibition

All above-mentioned data about *ERBB4* overexpression patterns due to *IKZF2-ERBB4* gene fusion led us to test tumour sensitivity to anti-HER4 inhibition in AC17 cells. For this purpose, we used different approaches: inhibition with chemical ERBB4 inhibitors used in clinical setting on different cell models and specific *ERBB4* knock outing. Indeed, we took the opportunity of *in vivo* and *in vitro* models we derived from the patient TumAC17: the PdxAC17 xenograft and the subsequent derived AC17 cell cultures.

We have selected afatinib, a pan ERBB-inhibitor known to be active against ERBB4 and approved for clinical use [Nakamura et al, 2016] as specific anti-ERBB4 did not yet exist. First *in vitro* experiments on tumour cells showed no effect of afatinib at low and moderate concentration but drastic cell toxicity at high concentrations from 5 to 10 µM (**Figure 4A-4B**). Such efficient concentrations are non-relevant for clinical application since pharmacokinetics studies have reported that plasma concentration of afatinib never reaches 100ng/ml *ie* 0.2µM [Wind et al, 2017]. This absence of effect on tumour cells were confirmed in *in vivo* assay in mice xenografted with PdxAC17 (**Figure 4C**). No statistical difference in tumour size was observed between control *versus* treated group (Mann-Whitney assay, *p*=0.4848). As tumour size does not always reflect tumour cell viability or proliferation capacity, we performed blinding histology analysis on tumours at the end of *in vivo* RTK inhibitor assay. Tumour cells displayed neither different mitosis index nor necrosis rate between control and afatinib group (**Figure 4D**, Mann-Whitney assay, *p*=0.2684 and 0.6840 respectively). Nevertheless, Western Blot analysis confirmed the blocking of ERBB4 after afatinib exposure (**Figure 4E, Supplemental Figure S2**) through the inhibition of ERBB4 phosphorylation. Examination of activation of the downstream signaling PI3K/AKT and MAPK/ERK pathways, showed decrease of MEK phosphorylation at low concentration and in a less extent decrease of ERK phosphorylation, while AKT pathway was not impacted by afatinib with sustained phosphorylation of AKT and S6. Experiments with a second pan-ERBB inhibitor lapatinib targeting ERBB4 [Popović et al, 2024] led to similar results: no cell growth inhibition at therapeutic doses [Ohgami et al, 2017] (**Figure 4F**), no inhibition of AKT pathway while level of MEK phosphorylation was decreased with lapatinib incubation (**Figure 4G**, **Supplemental Figure S2**). The phenomenon of crosstalk between PI3K/AKT and MAPK/ERK pathways (*i.e.* inhibition of the PI3K/AKT pathway leading to activation of the MAP kinase ERK and, conversely, inhibition of the MAPK/ERK pathway increasing phosphorylated AKT levels) largely reported in therapeutic resistance [Turke et al, 2012] could explain the lack of efficacy of ERBB4 inhibition. In addition to these 2 downstream pathways, ERBB4 is also known as STAT5A transcriptional activator by regulating STAT5A phosphorylation, notably through its intracellular domain (4ICD) released after gamma-secretase cleavage [Williams et al, 2004]. Incubation of AC17 cancer cells with low doses of afatinib (0.1 µm, close to clinical concentration) is poorly efficient to decrease expression of *BCL2*, *BCLXS*, *BCLXL*, *SOCS3*, *CISH*, *CCND1* and *MYC* genes (**Supplemental Figure S2)** selected as well-known STAT5 target genes [Halim et al, 2020]. Likewise, more specific inhibition of ERBB4 using siRNA approach (**Figure 4H**) did not lead to efficient cell growth decrease.

**Figure 4.**
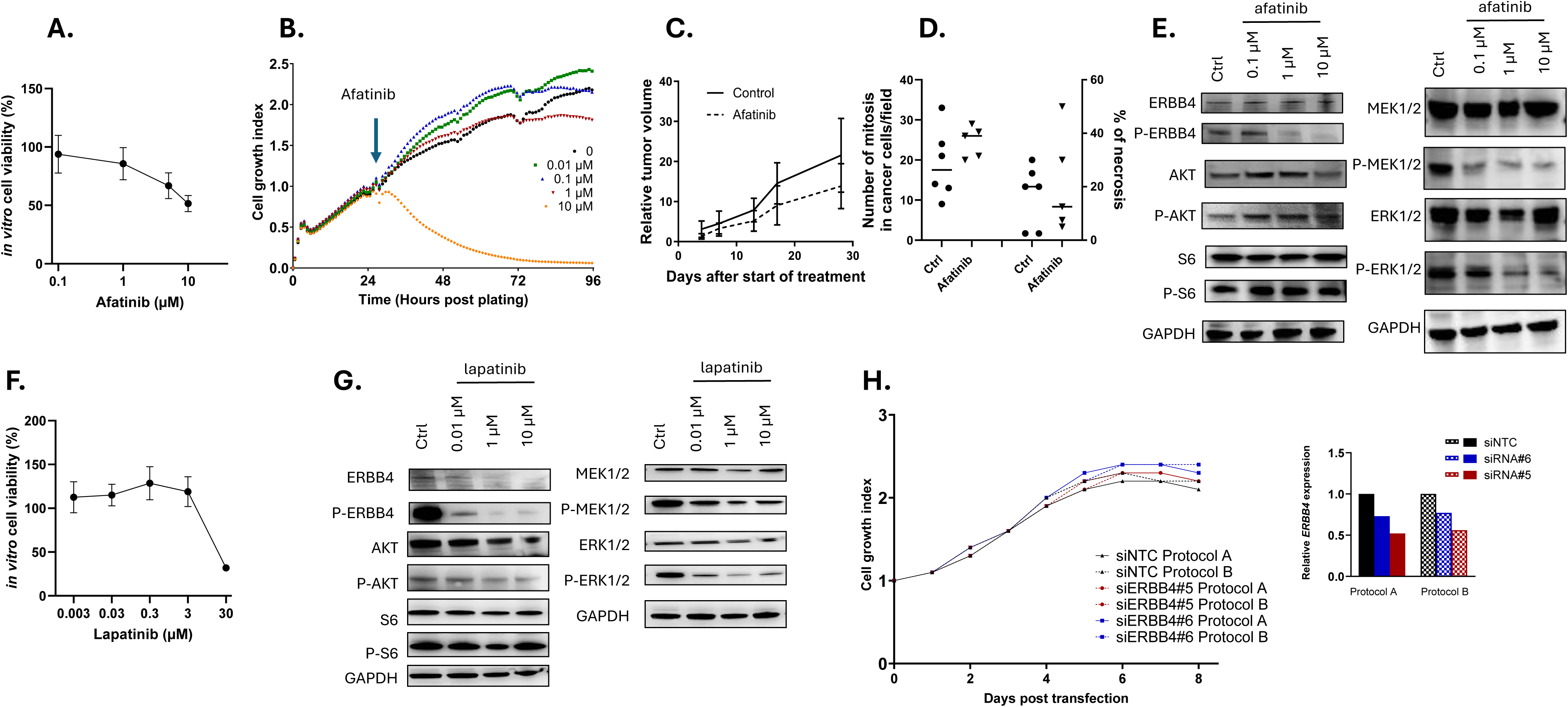
ERBB4 blocking does not lead to efficient cell growth inhibition in AC17 models. Viability assays of AC17 cells incubated with afatinib for 72h using WST (**A**) and xCELLigence (**B**). Nude mice have been subcutaneously xenografted with PdxAC17 for oral treatment by afatanib: (**C**) Tumor growth curve and (**D**) blind histological analysis at the end of the experiment. **E.** Western blot analysis of ERBB4 downstream signaling pathways in AC17 cells incubated with afatinib for 24h (a representative of 2 experiments is shown). **F.** Viability assays of AC17 cells incubated with lapatinib for 72h using WST. **G.** The expression levels of total and phosphorylated proteins were determined by Western blot analysis in cells incubated with lapatinib for 24h (one experiment). **H.** Lack of effect of siRNA-mediated ERBB4 knockdown on AC17 cell viability. AC17 cells were transfected with non-targeting (siNTC) or targeting *ERBB4* (siERBB4, number#5 and#6) siRNAS (a representative of 3 experiments is shown). *Left*: Cell viability was evaluated using Incucyte method. *Right*: qRT-PCR showing decrease of *ERBB4* mRNA.

Finally, we reported here the presence of *IKZF2-ERBB4* gene fusion one more time, but this time in ASCC, accompanied with *ERBB4* overexpression. RTK gene fusions are largely documented as important class of cancer-driving event with therapeutic and diagnostic value. Importantly, this *IKZF2-ERBB4* fusion has been already identified in other 3 solid tumour types, motivating the pancancer targetable potential of this rare gene fusion. However, according to the AACR Project GENIE Consortium [AACR Project GENIE Consortium, 2017], *ERBB4* is altered in 2.85% of all cancers making this RTK an attractive therapeutic target. Nevertheless, we showed that anti-ERBB4 blocking did not lead to decrease *in vitro* and *in vivo* cell growth, underlying the importance of combination therapies to overlap tumour resistance.

## Acknowledgements

We thank the In vivo experiment platform for mouse housing and the Department of Pathology, Institut Curie for their assistance in specimen collection and DNA/RNA preparation.

## Author contributions

Conceptualization: I.B and V.D-M. Methodology: S.R, A.S, J.L, A.B, S.V, R.E-B, E.E-A, P.M, C.N, A.L, W.C. Supervision and Writing—original draft: I.B and V.D-M. Writing—review & editing: S.R, A.S, S.V, R.E-B, I.B and V.D-M. The authors have neither financial disclosure or nor conflict of interest to declare.

## Data availability

The datasets used during the current study are available from the corresponding author on reasonable request.

## SUPPLEMENTARY INFORMATION

Supplemental Figure S1. *In situ* hybridization showing the expression of gene fusion transcript in PdxAC17 tumour cells.

Supplemental Figure S2. Pharmacological blocking of ERBB4 leads mainly to the inhibition of MEK downstream signaling pathway while AKT and STAT5 pathways are less affected.

Supplemental Table S1. Clinico-pathological features of ASCC tumour samples.

Supplemental Table S2. Sequences of primers.

Supplemental Table S3. Selection of genes encoding Receptor Tyrosine Kinases for potential targeted therapies.

## Notes

### Competing Interest Statement

The authors have declared no competing interest.

