## Supplementary material for "IKZF2-ERBB4 gene fusion leads to *ERBB4* overexpression in Anal Squamous Cell Carcinoma but with no benefit for ERBB4 inhibition": Suppl Figures S1&S2 Suppl Tables S1 S2 & S3

<u>Suppl Figure S1</u>. ***In situ* hybridization showing the expression of gene fusion transcript in PdxAC17 tumour tissue**.

**A.** *IKZF2-ERBB4* fusion transcript is stained in green while no red staining is visible for WT *ERBB4* transcript. **B**. Positive Control with human probe *PPIB* (in red) and *POLR2A* (in green). Scale bar: 50 µm.

**A.**

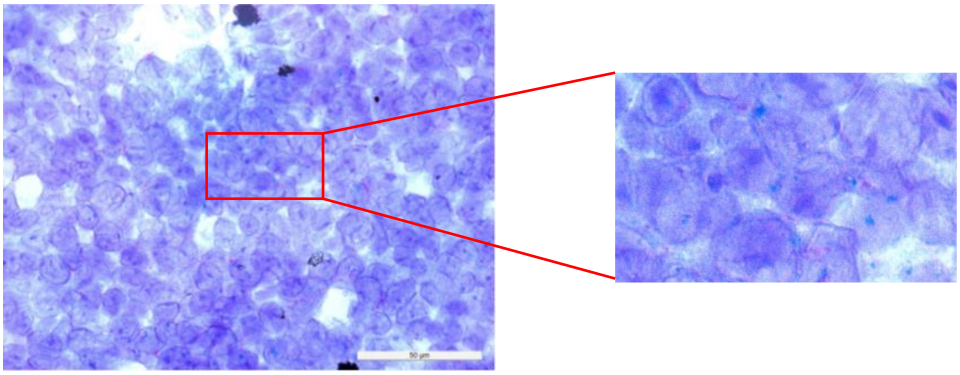

**B.**

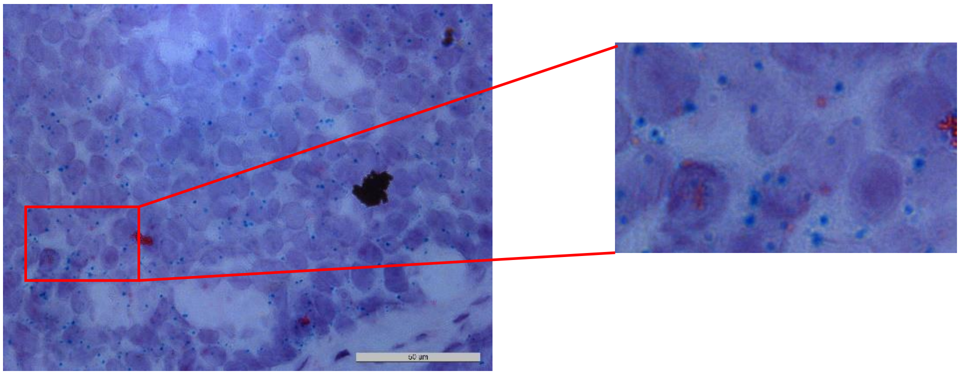

**Suppl Figure S2. Pharmacological blocking of ERBB4 leads mainly to the inhibition of MEK downstream signaling pathway while AKT and STAT5 pathways are less affected.**

**A.** Quantification of P-ERBB4, ERBB4, P-AKT, AKT, P-S6, S6, P-MEK1/2, MEK1/2, P-ERK1/2, ERK1/2 was performed by the Multi Gauge software and normalized on GAPDH expression. The ratio of Phosphorylated protein/Non phosphorylated protein is shown in afatinib or lapatinib 24h-treated AC17 cells. *Left*: Analysis of 2 WB with protein extracts from afatinib treated AC17 cells. *Right:* One WB with lapatinib treated AC17 cells. **B.** Genes have been selected for their induced expression through STAT5 activation. Gene expression results are expressed as N-fold differences in target gene expression relative to the TBP house-keeping gene.

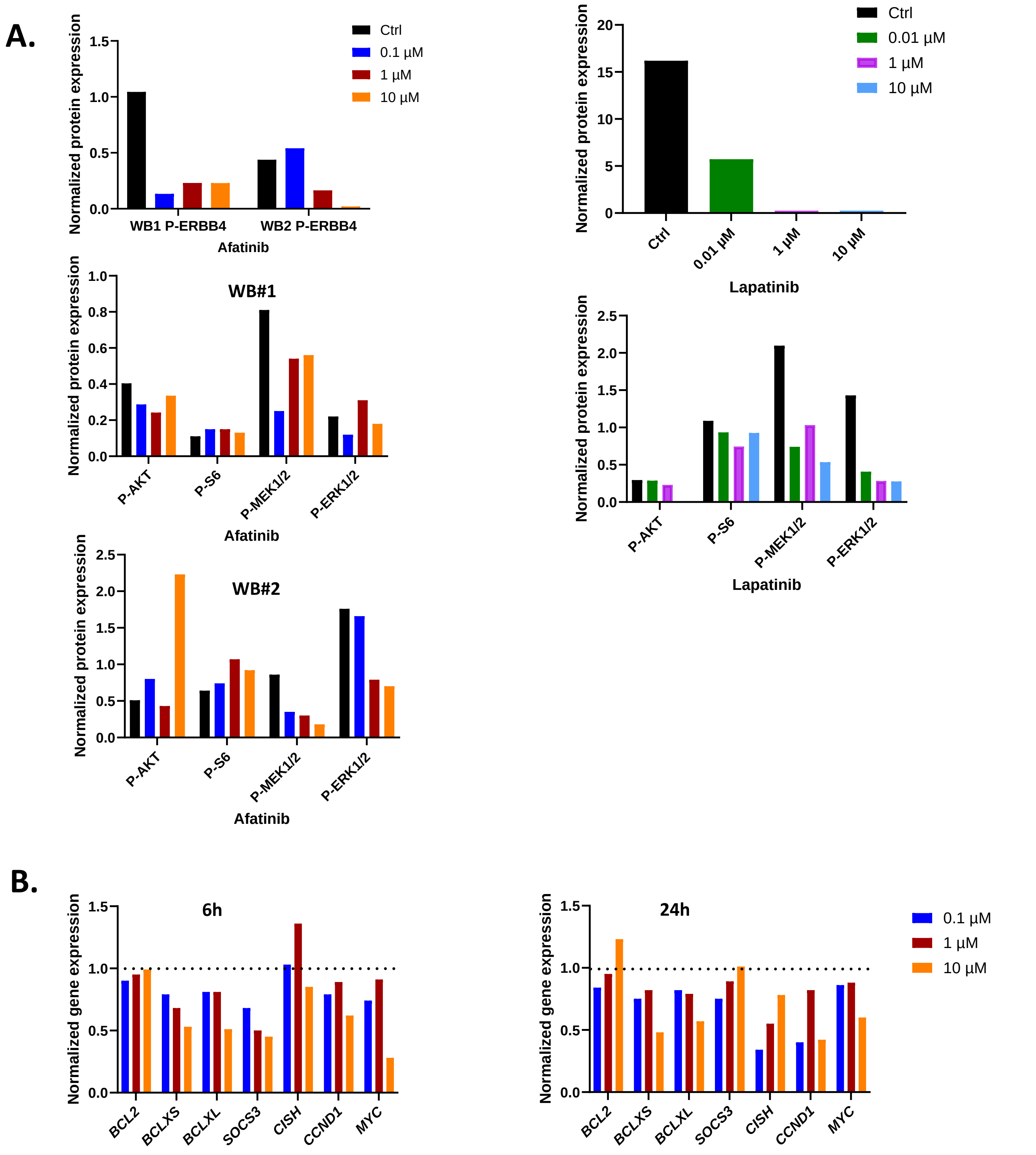

Richon et al.

**<u>Supplemental Table S1.</u> Clinico-pathological features of ASCC tumour samples.**

| **Characteristics** | | **Patients n=40** | |
|---|---|---|---|
| **Gender** | Female | 30 | 75% |
| | Male | 10 | 25% |
| **Tumour& metastasis site** | Anus | 35 | 88% |
| | Lymph node | 2 | 5% |
| | Liver | 2 | 5% |
| | Other | 1 | 3% |
| **Type of tumours** | Naive | 16 | 40% |
| | Relapse (after initial RT or RCT) | 24 | 60% |
| **Tumour differentiation** | Poor | 5 | 13% |
| | Moderate/Well | 35 | 88% |
| **HPV status** | HPV positive | 37 | 93% |
| | genotype 16 | 35 | 88% |
| | other genotypes (6-11/18) | 2 (1/1) | 5% |
| | HPV negative | 3 | 7.5% |
| **Concomitant HIV infection** | Yes | 3 | 7.5% |
| | No | 37 | 92.5% |

**<u>Supplemental Table S2</u>. Sequences of primers. A.** Primers used for quantitative RT-PCR for the 18 RTK and TBP control genes. **B.** Primers for gene fusion analysis. The 3 pairs have been used to demonstrate the presence of the fusion transcript and the *IKZF2/ERBB4* pair for amplification for PCR product sequenced with Sanger method.

## A.

| Gene | Upper sequence | Lower sequence | Amplicon size (pb) |
|---|---|---|---|
| ***ALK*** | 5' CGG AGG ATA TAT AGG CGG CAA T 3' | 5' ATG CCC AGT GGA CTG ATG AAG GA 3' | 90 |
| ***EGFR*** | 5' GGA GAA CTG CCA GAA ACT GAC C 3' | 5' GCC TGC AGC ACA CTG GTT G 3' | 106 |
| ***ERBB2*** | 5' AGC CGC GAG CAC CCA AGT 3' | 5' TTG GTG GGC AGG TAG GTG AGT T 3' | 147 |
| ***ERBB3*** | 5' GTCTGTGTGACCCACTGCAACT 3' | 5' GGGTGGCAGGAGAAGCATT 3' | 80 |
| ***ERBB4*** | 5' GGC TGC TGA GTT TTC AAG GAT G 3' | 5' GCT TCA TAC GAT CAT CAC CCT GA 3' | 74 |
| ***FGFR1*** | 5' GAA TTG GAG GCT ACA AGG TCC GTT 3' | 5' GGT TGA TGC TGC CGT ACT CAT TCT 3' | 118 |
| ***FGFR2*** | 5' GGC CGT GAA GAT GTT GAA AGA TG 3' | 5' CCT GTG TGC AGG CTC CAA GAA 3' | 128 |
| ***FGFR3*** | 5' GGCTGAAGAACGGCAGGGAGT 3' | 5' CTGCTGATGCCGCAGCTTGA 3' | 68 |
| ***IGF1R*** | 5' CCAAGGCCTGAAAACTCCATCT 3' | 5' ACACATTCTCGCTGATCCTCAACT3' | 116 |
| ***MET*** | 5' ATG GGT CAA TTC AGC GAA GTC C 3' | 5' GAT CGA GAA ACC ACA ACC TGC AT 3' | 119 |
| ***KIT*** | 5' TCC TCG CCT CCA AGA ATT GTA T 3' | 5' CTT GAT GTC TCT GGC TAG ACC AAA 3' | 111 |
| ***RET*** | 5' GCC ACC GAC CAG CAG ACC T 3' | 5' GCC TCC TCG GCC ACA TAT GA 3' | 80 |
| ***CSF1R*** | 5' GATGGGTGGCAGGAAGGTGA 3' | 5' GGGCCCTGGGATGACTTTCT 3' | 67 |
| ***PDGFRA*** | 5' CAT TTA CAT CTA TGT GCC AGA CCC A 3' | 5' ATG GCA GAA TCA TCA TCC TCC AC 3' | 93 |
| ***PDGFRB*** | 5' CCC CAG TGC CGA GTT AGA AGA C 3' | 5' GCA CGT AGC CGC TCT CAA CC 3' | 116 |
| ***VEGFR1*** | 5' ATC ATT CCG AAG CAA GGT GTG AC 3' | 5' TCC TTC TAT TAT TGC CAT GCG CT 3' | 122 |
| ***VEGFR2*** | 5' TGG GAA CCG GAA CCT CAC TAT C 3' | 5' GTC TTT TCC TGG GCA CCT TCT ATT 3' | 132 |
| ***VEGFR3*** | 5' GTC ACG CTG CGC TCG CAA A 3' | 5' GTG CAG CAG TGG CGT GGA CA 5' | 99 |
| ***TBP*** | 5' TGC ACA GGA GCC AAG AGT GAA 3' | 5' CAC ATC ACA GCT CCC CAC CA 3' | 112 |

## B.

| Primers name | Sequences | Amplicon size (pb) |
|---|---|---|
| ***ERBB4_exon 1-2*** | 5’ AGGACTTTGGGTCTGGGTGA 3’ | 230 |
| ***EERBB4_exon 1-2*** | 5’ CTCGAACAGACCGCAGGAA 3’ | |
| ***ERBB4_exon 4-5*** | 5’ ATTCCTTTGTTATGCAGACACCA 3’ | 130 |
| ***ERBB4_exon 4-5*** | 5’ CCAGCAACGGCCAGTACAG 3’ | |
| ***IKZF2/ERBB4*** | 5’ TGACTATGGAAACAGAGGCTATTG 3’ | ≈500 |
| ***IKZF2/ERBB4*** | 5’ CCACCATTTAGGATTTCTGTCAA 3’ | |

**Supplemental Table S3. Selection of genes encoding Receptor Tyrosine Kinases for potential targeted therapies.**

| Gene | Names of genes | Chromosome location | Genbank accession number | Examples of targeted therapies |
|---|---|---|---|---|
| ***EGFR*** | epidermal growth factor receptor | 7p12 | NM_005228.3 | cetuximab, gefitinib, erlotinib, lapatinib, afatinib, neratinib, panitumumab, osimertinib, |
| ***ERBB2*** | erb-b2 receptor tyrosine kinase 2 | 17q12 | NM_004448.3 | trastuzumab, lapatinib, afatinib, neratinib |
| ***ERBB3*** | erb-b2 receptor tyrosine kinase 3 | 12q13 | NM_001982.3 | pertuzumab, MM121 |
| ***ERBB4*** | erb-b2 receptor tyrosine kinase 4 | 2q33.3-q34 | NM_005235.2 | lapatinib, afatinib, neratinib |
| ***FGFR1*** | fibroblast growth factor receptor 1 | 8p11.23- | NM_015850.3 | vandetanib, PD173074, erdafitinib |
| ***FGFR2*** | fibroblast growth factor receptor 2 | 10q26 | NM_000141.4 | vandetanib, PD173075, erdafitinib |
| ***FGFR3*** | fibroblast growth factor receptor 3 | 4p16.3 | NM_000142.4 | vandetanib, PD173076, dovitinib, erdafitinib |
| ***IGF1R*** | insulin-like growth factor 1 receptor | 15q26.3 | NM_000875.4 | linsitinib, dalotuzumab, figitumumab |
| ***MET*** | MET proto-oncogene, receptor tyrosine | 7q31 | NM_000245.2 | crizotinib, foretinib, capmatinib, tepotinib, savolitinib, onartuzumab |
| ***ALK*** | anaplastic lymphoma receptor tyrosine | 2p23 | NM_004304.4 | Crizotinib, alectinib, ceretinib, lorlatinib |
| ***PDGFRA*** | platelet-derived growth factor receptor, | 4q12 | NM_006206.4 | imatinib, sunitinib, sorafenib, pazopanib, vatalanib, tandutinib, |
| ***PDGFRB*** | platelet-derived growth factor receptor, beta | 5q33.1 | NM_002609.3 | nilotinib, |
| ***VEGFR1*** | fms-related tyrosine kinase 1 | 13q12 | NM_002019.4 | sunitinib, sorafenib, pazopanib, vatalanib, , cediranib, vandetanib |
| ***VEGFR2*** | kinase insert domain receptor | 4q11-q12 | NM_002253.2 | sunitinib, sorafenib, pazopanib, vatalanib, apatinib, brivanib, , |
| ***VEGFR3*** | fms-related tyrosine kinase 4 | 5q35.3 | NM_002020.4 | sunitinib, sorafenib, pazopanib, vatalanib, , cediranib, vandetanib |
| ***KIT*** | v-kit Hardy-Zuckerman 4 feline sarcoma viral | 4q12 | NM_021099.3 | imatinib, sunitinib, sorafenib, pazopanib, vatalanib, tandutinib, |
| ***CSF1R*** | colony stimulating factor 1 receptor | 5q32 | NM_005211.3 | sunitinib, vatalanib |
| ***RET*** | ret proto-oncogene | 10q11.2 | NM_020630.4 | sunitinib, motesanib |
